# Barriers to coexistence limit the poleward range of a globally-distributed plant

**DOI:** 10.1101/2020.02.24.946574

**Authors:** David W. Armitage, Stuart E. Jones

## Abstract

Species’ poleward ranges are thought to be primarily limited by climatic constraints rather than biotic interactions such as competition. However, theory suggests that a species’ tolerance to competition is reduced in harsh environments, such as at the extremes of its climatic niche. This implies that under certain conditions, interspecific competition near species’ range margins can prevent the establishment of populations into otherwise tolerable environments and results in geographic distributions being shaped by the interaction of climate and competition. We test this prediction by challenging an experimentally-parameterized mechanistic competition model to predict the poleward range boundaries of two widely co-occurring and ecologically-similar aquatic duckweed plants. We show that simple, mechanistic ecological niche models which include competition and thermal response terms best predict the northern range limits of our study species, outperforming competition-free mechanistic models and matching the predictive ability of popular statistical niche models fit to occurrence records. Next, using the theoretical framework of modern coexistence theory, we show that relative nonlinearity in competitors’ responses to temperature fluctuations maintains coexistence at the subordinate competitor’s poleward range boundary, highlighting the importance of this underappreciated fluctuation-dependent coexistence mechanism. Our results demonstrate the predictive utility of mechanistic niche models and support a more nuanced, interactive role of climate and species interactions in determining range boundaries, which may help explain the conflicting results from previous tests of classic range limit theory and contribute to a more mechanistic understanding of range dynamics under global change.

## Introduction

Ecology has long clung to Darwin’s hypothesis [1] that species’ range limits are determined by biotic interactions toward the equator and by climatic harshness toward the poles [2, 3, 4]. This hypothesis assumes that the number of other species or individuals with which a species interacts increases towards the equator, eventually reaching a latitude where the effects of competition, predation, or disease curb further expansion. Because species richness and population densities often decline toward the poles, environmental stress is thought to have primacy over interspecific interactions in determining species’ poleward boundaries. However, empirical support for this century-old hypothesis — variously called *stress gradient hypothesis* or the *species interactions–abiotic stress hypothesis* (SIASH) [5]— remains mixed [6, 7, 8, 9, 10].

These hypotheses commonly posit that the *per capita* magnitude [11, 5] or relative frequency [12, 13] of negative interactions, such as competition, should decrease with environmental stress resulting in weaker competitive regulation of populations in harsher environments. While this prediction often holds (reviewed in [5]), a population’s *tolerance* of competition can also decrease in harshening environments, where even weak competition can drive an already-low *per capita* growth rate, *r* (= *dN/Ndt*, where *N* is population size), below zero [14, 15, 16, 17, 18, 19]. The relative importance of competition or other negative interactions in shaping range margins may therefore hinge on a balance between the overall intensity of competition experienced by a marginal population and the extent to which competition suppresses its environmentally-determined *per capita* growth rate.

Assuming that species’ geographic distributions are manifestations of their ecological niches [20, 21, 22], it becomes possible to study factors shaping these distributions using the quantitative tools of population ecology [23, 24]. A species’ biotically-reduced or realized niche can be defined by the combined abiotic and biotic states over which its intrinsic *per capita* growth rate, *r*, is greater than zero, indicating persistence is possible. This rate can be expressed as a function, *r*(*E, C*), of the both the local abiotic environment (*E*) and the effects of biotic interactions such as competition (*C*) [25]. For a species that does not interact with any others nor experience dispersal limitation, its geographic range is limited solely by its growth response to the abiotic environment (i.e., its fundamental niche) [20] (Fig. 1A). Under an SIASH scenario, stressful abiotic conditions at a species’ poleward margin will cause a zero net-growth boundary, beyond which persistence cannot be sustained 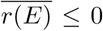, where the overbar indicates a long-term time average over environmental fluctuations). However, if 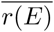 cannot sufficiently predict an observed range boundary, then we are left to consider alternative range-limiting mechanisms such as competition, predation, and mutualism — the effects of which can modify the niche in a variety of ways [22, 24] (Figs. 1B and 1C).

**Fig. 1.**
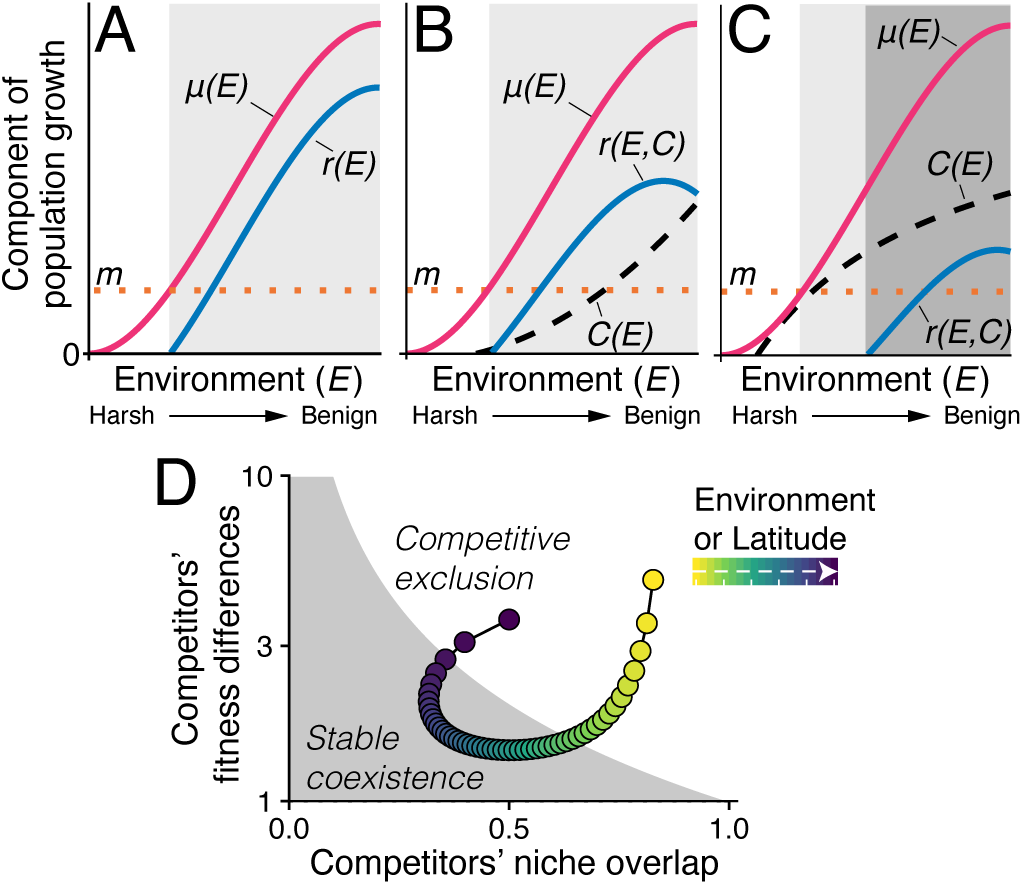
A species’ intrinsic growth rate, *r* (= *dN/Ndt*), can be expressed as a function of its birth rate, *μ*, the impacts of competition from other species, *C*, and a mortality term, *m*, such that *r* = *μ* − *m* − *C*. We can define a species’ niche breadth as the conditions where *r >* 0 [24]. (A) In the absence of interspecific interactions, the gray area shows the fundamental niche for a species with a monotonic growth response across an environmental gradient *E*. For simplicity, we assume mortality *m* is constant over *E*. (B) Consistent with an SIASH pattern [5], including a competition term *C*(*E*) that decreases in harshening environments suppresses the *per capita* growth but does not affect the niche limit. (C) However, slightly adjusting the functional form of *C*(*E*) causes the niche to be truncated by competition in harsh environments by negating the positive effects of *μ*(*E*) [15]. The darker gray region shows the new biotically-limited niche breadth. (D) Range boundaries set by competitive interactions are termed *coexistence boundaries*, and occur where a species’ long-term, low-density growth rate in the presence of resident competitors switches sign. This growth rate is determined by a combination of fitness differences and niche overlap with competitors — both of which covary over environmental or latitudinal gradients.

**Fig. 2.**
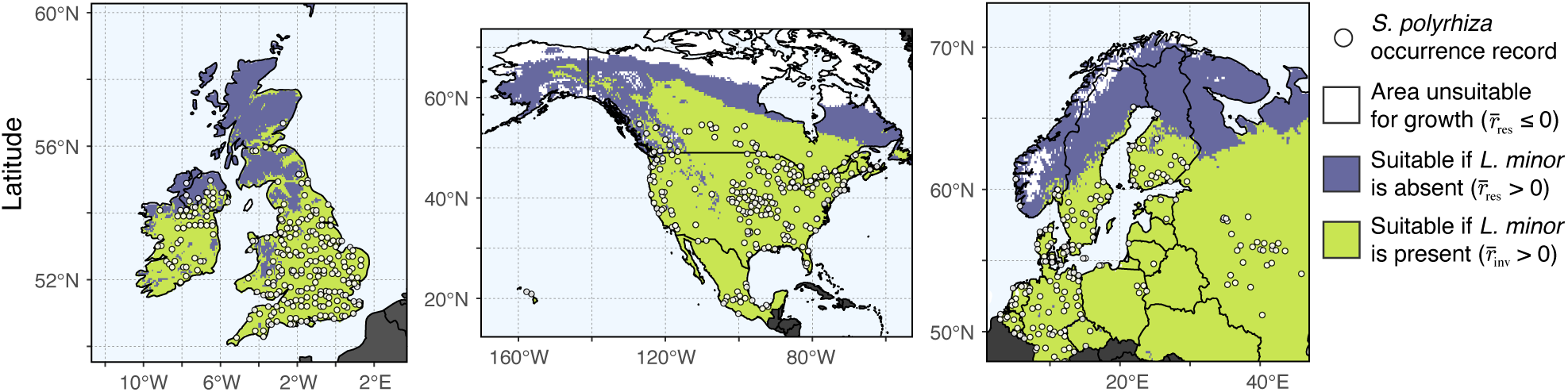
Range predictions for *S. polyrhiza* from the competition model (eq. 1) projected across geographic space. Shading denotes areas of predicted population persistence where long-term low-density growth rates in the absence 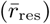 or presence 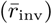 of resident competitor *L. minor* is greater than zero.

Quantifying the joint, interactive effects of the abiotic environment and competition on species’ growth rates is challenging, but can be accomplished using the tools of modern coexistence theory (MCT) [26]. If a species’ latitudinal range margin is limited by competition, it is incapable of stably coexisting with resident competitor species at and beyond the latitude where its long-term *invasion growth rate*, 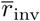, switches sign from positive to negative. This growth rate quantifies a species’ ability to invade and therefore coexist with a community of resident competitors, and can be partitioned into the relative contributions of various coexistence mechanisms [27]. These mechanisms reduce species’ niche overlap and growth advantages to prevent competitive exclusion, and are defined by their degree of dependence on fluctuations in environmental and competitive factors [26]. In a geographic context, coexistence outcomes are predicted to vary over space if the environment covaries with latitude, leading to the prediction that competition-limited species ranges manifest where the joint effects of the environment and competition prohibit coexistence (Fig. 1D) [28]. Bringing the MCT framework to bear on niche models permits an explicit quantification of mechanisms causing competitor-limited range margins, though this has yet to be attempted using real distributional data [24, 29, 30].

Here, we ask whether a dynamic, mechanistic niche model [31] can accurately predict the poleward range margins of two of the most broadly-distributed plants on Earth — the duckweeds *Lemna minor* and *Spirodela polyrhiza*. These minute floating plants widely coexist in fresh waters across N. America, Eurasia, Africa, and Australia (*SI Appendix*, Figs. S1 and S2), though competition limits their stable coexistence under certain conditions [32, 33]. Our objective was to test whether species’ laboratory-measured growth responses to temperature and competition can be used to accurately predict their poleward range limits. This approach has successfully predicted species distributions over various niche dimensions [34, 35], but has yet to be used for interrogating the determinants of geographic range margins. As our niche model explicitly accounts for the effects of competition on invasion growth rates, we ask whether predicted range limits are more accurate in the presence or absence of interspecific competition. We then partition these invasion growth rates into their constituent coexistence mechanisms to identify how these mechanisms vary across space and whether they act to maintain competitors’ poleward range boundaries.

## Materials and Methods

### Mechanistic Niche Model

Detailed methods are described in *SI Appendix*. We use a previously-developed stage-structured differential equation model describing the population growth rates of our two focal competitor species. The process for developing and fitting this model is detailed in [33]. The model describes the temporal dynamics of the species’ (*j*) vegetative and dormant (turion) forms (*N*_*j*_ and *S*_*j*_, respectively) according to the equations

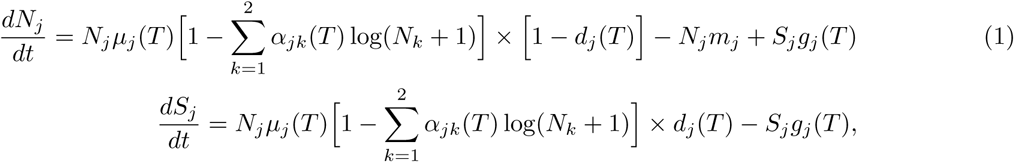

where *m*_*j*_ is a constant *per capita* mortality rate for each species. The maximum daily *per capita* growth rate, *μ*_*j*_(*T*), is a unimodal function of temperature following the expression

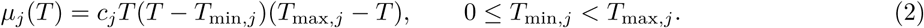

Here, the parameters *T*_min,*j*_ and *T*_max,*j*_ describe the minimum and maximum temperatures at which growth is possible, and *c*_*j*_ is a shape constant. Inter- and intraspecific competition were modeled using the temperature-dependent parameters *α*_*jk*_ (*k* = {1, 2}) described by the function

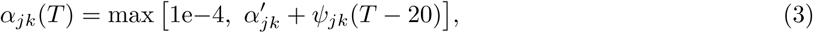

where 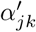 are competition coefficients measuring the proportional effect of *N*_*k*_ on the growth rate of species *j* at 20°C, and *ψ*_*jk*_ describe the change in the strength of competition with ambient temperature. We prevent the competition parameter from switching sign by assigning an arbitrarily low, positive value to instances where it would otherwise be less than or equal to zero. We use the logarithm of competitor density to describe each species’ concave-up density-dependent growth responses. Instantaneous temperature-dependent turion investment, *d*_*j*_, and germination, *g*_*j*_, fractions were modeled as logistic functions of temperature using the equations

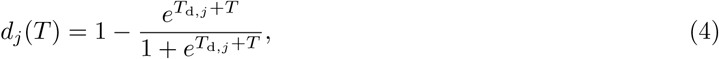

and

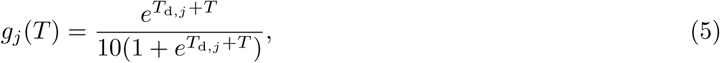

where *T*_d,*j*_ is the temperature at which turion production accounts for 50% of total new growth, and *T*_g,*j*_ is the temperature at which 50% of turions have germinated after 10 days. Model parameters were empirically estimated from replicated growth and competition assays conducted in environmental chambers spanning a range of ambient temperatures from 3°C to 37°C (*SI Appendix*, Fig. S3), and had acceptable predictive accuracy (*SI Appendix*, Fig. S4).

### Mechanistic Range Prediction

We used 2.5 arcmin-resolution climate data [36] to generate 10 years of temperature fluctuations for each grid cell. Using these time series, we simulated the population dynamics of each species to monoculture equilibria, saving at each time step its *per capita* growth rate, *r*_res_(*t*) (= (*N*_*j*_ + *S*_*j*_)^−1^ (*dN*_*j*_*/dt* + *dS*_*j*_*/dt*)). Here, the subscript ‘res’ indicates that the species is in its resident, monoculture state and so interspecific competition does not occur. After verifying 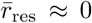 over the last year of the simulation, we saved the equilibrial abundances. We used each grid cell’s temperature and resident population time series to estimate the long-term average growth rates of an invading species 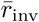. We set resident densities *N*_*k*_ at equilibrium and conspecific densities *N*_*j*_ at one, then calculated *r*_inv_(*t*) at each time step over the final year, time-averaging the resulting growth rates. We identified the set of cells where each species’ long-term low-density growth rates as residents 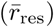 and invaders 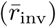 were greater than zero. The former set are range predictions derived from the fundamental niche, while the latter set, which includes the effects of competition, are range predictions derived from the fundamental niche. Coexistence is predicted where 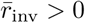 for both species in their invasion states [37].

### Statistical Range Prediction

Occurrence records of *S. polyrhiza* and *L. minor* used to fit our models were obtained from the Global Biodiversity Information Facility (GBIF) and Botanical Society of Britain & Ireland’s (BSBI) geo-referenced databases. We focused our analyses on three regions possessing an abundance of high-quality botanical records and encompassing our species’ northern latitudinal range limits. These regions include the United Kingdom and Ireland, North America (Mexico, US, and Canada), and Northern Continental Europe. Global records were accessed and downloaded from GBIF and BSBI, which were then quality-filtered [38].

We used the maximum entropy method (MaxEnt) to predict the distributions of *L. minor* and *S. polyrhiza* [39]. Models were fit to spatially-thinned occurrence records for each species in all three study regions, as well to the combined suite of point records across all study regions. Covariates included two sets of 2.5 arcmin bioclimatic variables. The first group of models, called MaxEnt_12_, used twelve bioclimatic covariates while the second group, called MaxEnt_2_, include only mean temperature and annual temperature amplitude — the same two covariates used in our mechanistic models’ predictions. MaxEnt models were fit using 4-fold cross validation across nested, geographically-independent checkerboard partitions [40] and across a range of regularization parameters [41]. Best-fit models were selected based on relative AIC rankings [42], 10% omission rate metrics [40], and by visual inspection of the results. Binary predictions of species’ ranges were made by thresholding the predicted occurrence probabilities, *p*_occ_, by 10% omission threshold, *τ*, to ensure at least 90% of location records are included within the range [43].

### Evaluating and Comparing Model Predictions

We used a beta regression with a logit link function [44] to assess the relationship between predicted invasion growth rates and MaxEnt occurrence probabilities. True poleward range limits limit was estimated from observation records by calculating each species’ 95% confidence intervals for each region’s latitudinal maxima. These maxima were estimated across longitudinal bins of various widths depending on the region. Niche model estimates were regressed against latitude to identify *x*-intercepts and associated and their associated inverse 95% confidence intervals [45] corresponding to latitudes at which 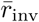 and *p*_occ_ − *τ* were zero (Fig. 3A).

**Fig. 3.**
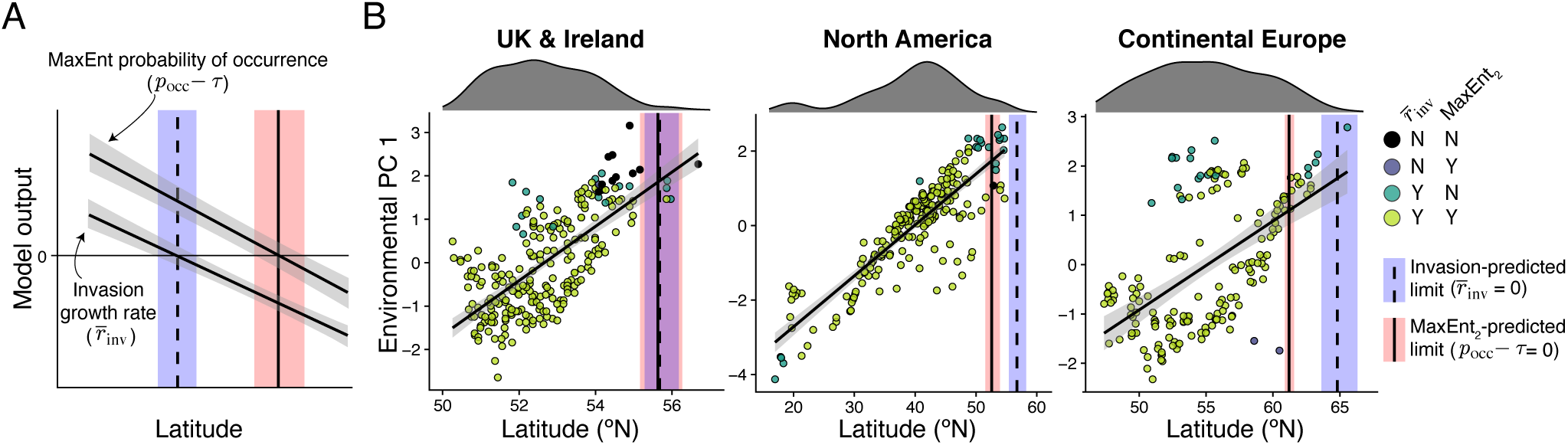
(A) Conceptual illustration of range limits for *S. polyrhiza* are predicted from niche model outputs. Model outputs were regressed against latitude and their *x*-intercepts (± 95% CI) were used to determine the predicted latitudinal limits. For MaxEnt models, outputs were the probability of occurrence *p*_occ_ minus the 10% omission threshold value (*τ*). For mechanistic models, we use the predicted long-term invasion growth rate 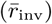. (B) We detected strong associations between latitude and aggregated environmental covariates (here, average annual temperature and its amplitude) in all three regions. Points denote *S. polyrhiza* observation records, colored by binary model classification results, and vertical lines show the estimated mean latitudinal limits (± 95% CI) for each model and region. Histograms above each plot show the latitudinal dispersion of occurrence records after spatial thinning.

### Quantifying Coexistence Mechanisms

We estimated the contributions of various fluctuation-dependent and fluctuation-independent mechanisms on the coexistence of *S. polyrhzia* and *L. minor* (*SI Appendix*) [27]. This method partitions differences between the long-term growth rate of an invading species, 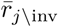 and that of a resident, 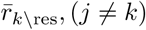 (which is approximately zero at equilibrium), using the equation

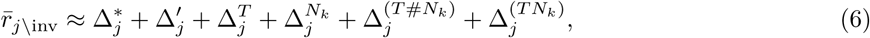

where 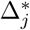 is the fluctuation-free growth rate, 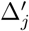 is the contribution of fluctuation-driven change in mean competitor density, 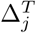 is the contribution of relative nonlinearity in temperature-growth responses, 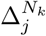 is the contribution of relative nonlinearity in responses to competitor densities, 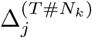 is the is the interaction between competitor density and temperature variability, and 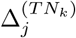 is the covariance between temperature and competition, which quantifies the temporal storage effect. These values were calculated across a range of average temperature and annual temperature variation to identify regions where particular coexistence promoting mechanisms operate. We overlaid global observation records for each species on these grids to identify (1) the environmental state-spaces where coexistence was predicted to break down relative to the observed environmental distribution of each species, and (2) which coexistence mechanisms contributed most strongly to shaping these coexistence margins.

## Results and Discussion

### Accounting for interspecific competition improves poleward range limit estimates

Using a mechanistic niche model to predict the invasion growth rates of *S. polyrhiza* in the absence 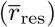 and presence 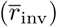 of interspecific competition, we found that range predictions made using 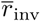 closely matched the distribution of observation records, while predictions from 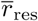 did not (Fig. 2, *SI Appendix*, Table. S5). We also encountered close correspondence between the observed latitudinal limits of *S. polyrhiza* and the competition model-predicted maximum latitude, particularly for the UK + Ireland and N. Europe regions (Fig. 3B-D, *SI Appendix*, Fig. S6 and Table S5).

Across all study regions, our mechanistic competition model performed as well as the best-fit statistical niche models in predicting latitudinal maxima. Though all models had high true positive rates, binomial omission tests indicated only MaxEnt models and mechanistic competition models yielded predictions significantly better than random, while mechanistic ENMs without competition terms terms did not (*SI Appendix*, Table. S5). In contrast to our results for *S. polyrhiza*, we encountered no differences in the predicted poleward extents of *L. minor* (the superior competitor) regardless of whether or not interspecific competition terms were included *SI Appendix*, Fig. S6).

These results highlight the effects of interspecific competition on species’ poleward ranges. While the overall strength of interspecific competition declined with temperature (all *ψ*_*jk*_ > 0), its negative *per capita* effects on *S. polyrhiza* growth rates outpaced reductions in its environmental response. This observation contradicts prevailing hypotheses concerning the primacy of abiotic factors in structuring poleward range limits. While our data are consistent with the expectation that the strength of competition declines in harsh environments, it challenges the notion that the relative *importance* of competition should also decrease compared to abiotic factors. Accordingly, our results align with theoretical and empirical results showing that growth rates become increasingly sensitive to the effects of competition in suboptimal environments [14, 15, 16, 17, 18, 19].

We encountered a significant, positive correspondence between statistical and mechanistic competition model outputs for both *S. polyrhiza* and *L. minor* (*SI Appendix*, Figs. S7 and S8). This result illustrates an important point concerning the utility of statistical niche models for estimating species’ environmental responses. In areas where a species’ intrinsic growth rate is regulated by a competitor, the pure environmental responses of the focal species cannot be statistically recovered. This is because environmental responses extracted from occurrence records are confounded with the latent effects of biotic interactions. While this issue has been recognized for some time [46, 47, 48], it is commonly neglected when projecting species’ ranges using statistical niche model outputs. And while newer multispecies niche models can begin to parse biotic and abiotic responses for similar species, we remain unable to do so when a species occurs in nested sympatry with its competitors, as is the case for *S. polyrhiza* and probably many other organisms.

### Thermal fluctuations maintain the coexistence boundary of *S. polyrhiza*

We asked how different coexistence mechanisms varied in strength across niche axes of temperature averages and temperature fluctuations. Partitioning the invasion growth rate, 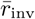, of *S. polyrhiza* into its constituent fluctuation-dependent and independent contributions, we found that its predicted coexistence boundary (the isocline where 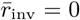) aligns very closely with the true distributional margin of its global occurrence records. This boundary is associated with a negative fluctuation-free growth rate Δ*, offset by a positive response to temperature fluctuations, Δ^*T*^ (Fig. 4). At the invasion boundary, Δ^*T*^ contributed positively to both species’ low-density growth rates while also reducing the growth rate differences that favor *L. minor* over *S. polyrhiza* (*SI Appendix*, Fig. S11). In contrast, we found no evidence that relative nonlinearity in competition Δ^*N*^ and the storage effect Δ^(^*TN*) contribute to maintaining the coexistence boundary (Fig. 4). The mechanisms operating near this boundary for *L. minor* are qualitatively similar, though the mean effects of temperature (Δ*) are stronger relative to those of temperature fluctuations (Δ^*T*^) (*SI Appendix*, Fig. S10).

**Fig. 4.**
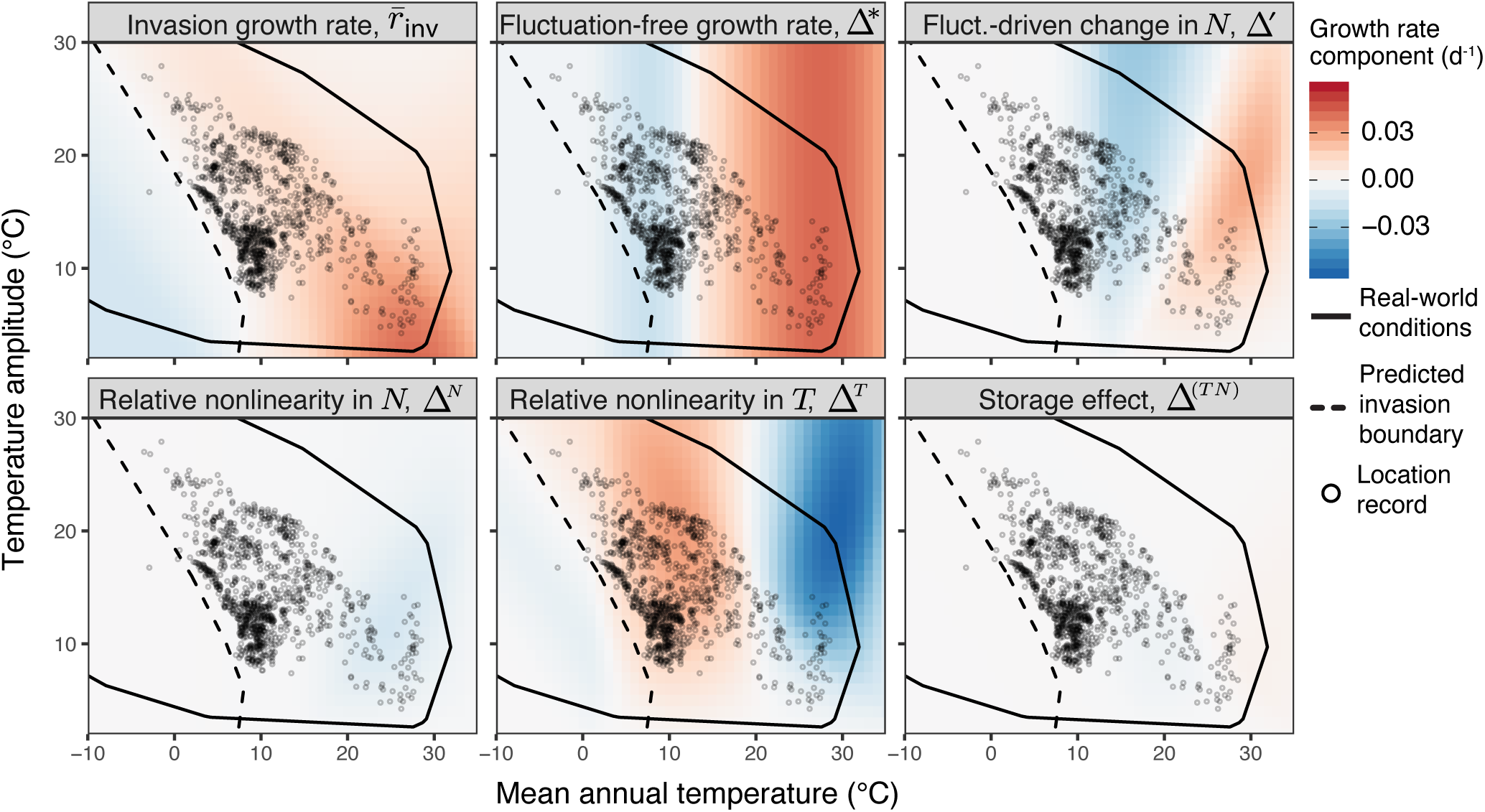
The upper-left panel displays the long-term invasion growth rate, 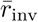, of *S. polyrhiza*, as a function of annual temperature averages and amplitudes. The dashed line denotes the model-predicted coexistence boundary (where 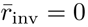), and the solid line encompasses the temperature regimes that occur across Earth. Points represent global occurrence records for *S. polyrhiza* projected onto their local temperature axes. The remainder of the panels quantify the contributions of the partitioned coexistence mechanisms from equation 6. Note the crucial role of relative nonlinearity in temperature responses, Δ^*T*^, in maintaining coexistence near the boundary.

That global occurrence records for *S. polyrhiza* overwhelmingly occur in regions where Δ* < 0 and Δ^*T*^ > 0 suggests that over much of the known global range of this species, temperature fluctuations, mediated through differences in competitors’ nonlinear thermal responses, are critical for maintaining positive *per capita* growth rates. Previous studies have demonstrated the important roles of fluctuation-dependent coexistence mechanisms such as relative nonlinearity in competition (Δ^*N*^) [49] and the temporal storage effect (Δ^(^*TN*)) [50]. However, the role of relative nonlinearity in environmental responses (Δ^*T*^) is one that, until recently, was not explicitly accounted for in MCT, yet is clearly important in cases where competitors’ environmental responses are nonlinear and nonidentical [27]. These results further demonstrate the degree to which coexistence mechanisms can rapidly shift across environments, implying that locally-measured coexistence-promoting mechanisms cannot be assumed uniform across a species’ range.

Despite the concordance between the observed and model-predicted ranges, our mechanistic niche modeling approach is not without limitations. First, we did not account for the effects of pathogens or predators in our model and assumed their effects were negligible. While anecdotal evidence supports this assumption [51], future studies are needed to quantify the differential effects of natural enemies on duckweed growth rates. Second, our focal species often occur in complex communities of plants and phytoplankton, and so our two-species model may underestimate the strength of interspecific regulation. Though small floating plants do co-occur with our focal species at lower latitudes, they are absent or very rare near and above the poleward limit of *S. polyrhiza*. Submerged macrophytes, on the other hand, are very common at high latitudes, but in most cases do not appear to prevent duckweed from successfully invading [52]. Third, while duckweed strains exhibit phenotypic variation in their competitive and environmental responses [53, 32], our mechanistic niche models use mean values extracted from regression parameters. While identifying how variation around these means affects geographic range predictions is beyond the scope of this study, our mechanistic model can be expanded to investigate these effects. Finally, while setting the MaxEnt binary prediction threshold *τ* to its 10% omission value is justifiable, other sensible choices would result in different poleward limit predictions. However, our choice of the 10% omission threshold is supported by many beta regression fits passing near where 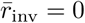 and *p*_occ_ −*τ* = 0 (*SI Appendix*, Figs. S7 and S8). When models are compared in this way, the mechanistic model’s natural threshold of zero net growth can guide users toward biologically-meaningful choices of *τ* in their statistical models.

Our study responds to calls for a more mechanistic, population ecology-based integration of niche theory with biogeography [21, 23, 54, 24, 48]. Using an experimentally-parameterized dynamic competition model, we are among the first to demonstrate, contrary to prevailing expectations, that interspecific competition can plausibly explain a species’ poleward range limit. We note, however, that this was only the case for our inferior competitor, *S. polyrhiza*, and the general extent to which competition influences species’ range boundaries remains an open question. It is likely that the importance of competition and other biotic interactions for range delimitation depends on how these factors covary across abiotic environmental gradients and influence the abiotically-determined fundamental niche (Fig.1). Quantifying such interactions will be a productive step toward understanding how and why communities and the coexistence mechanisms that maintain them vary over space [24, 29, 30].

From a methodological perspective, our results showcase the utility of mechanistic niche models for accurate range prediction without the need for occurrence records. With appropriate data, this framework could be employed to predict the effects of novel competitors encountered by species during climate-driven range shifts or following (re)introductions. It may also be valuable for predicting the distributions of species having few or no occurrence records using *ex situ* measurements of their environmental responses. Looking forward, we contend that geographically partitioning mechanistic niche models into their constituent coexistence mechanisms can fundamentally enhance our understanding of how populations and communities vary over space, potentially leading to strategies for promoting species coexistence in restoration, conservation, and relocation projects.

