## Supplementary material for "Barriers to coexistence limit the poleward range of a globally-distributed plant": SI Appendix

### Supporting Information Text

**Study Species.** The floating aquatic duckweeds *Spirodela polyrhiza* and *Lemna minor* are some of the smallest and most widely-distributed plants on earth, occupying all continents except Antarctica (1). The species have similar habitat preferences and frequently co-occur at both local (Fig. S1) and regional (Fig. S2) scales. While the ranges of these species show significant overlap, lake surveys have revealed that *L. minor* is the more abundant taxon in mixed *Spirodela-Lemna* communities. When present, *S. polyrhiza* is almost always encountered within larger crops of *L. minor* or the minute duckweed *Wolffia sp.*, whereas *L. minor* is more frequently encountered in dense monocultures (2). Dispersal occurs primarily via aquatic birds (3), though the poleward edges of both species' ranges terminate short of the the dispersal capabilities of migratory waterfowl and availability of freshwater habitats. The species exhibit different responses to cold temperatures, with *S. polyrhiza* producing dormant turion fronds and *L. minor* remaining viable as sunken vegetative fronds (4). Further, *S. polyrhiza* grows slightly better at higher temperatures, possibly explaining its higher prevalence in the tropics than *L. minor*, which grow better in cooler climates (1, 2).

**Estimating model parameters.** We measured low-density growth rates of *S. polyrhiza* and *L. minor* in temperature-controlled growth chambers as previously described (2). Briefly, axenic duckweed strains were grown in 100 mL flasks containing a chemically-defined medium approximating the natural mesotrophic pond waters from which they were collected (5). Growth assays were carried out for 28 days at static temperatures ($T$) ranging from 3°C to 37°C. Temperature-dependent low-density growth rates, $\mu_j(T)$ (day$^{-1}$), were measured from replicated monocultures inoculated with initial densities of 3 to 5 vegetative fronds. Growth at these densities was assumed exponential and estimated using the formula $\mu = [\log(N_t/N_0)]t^{-1}$. These values were used to fit thermal growth curves for each species, the best-fitting of which is shown as eq. 2 (6) (Fig. S2). Inter- and intraspecific competitive responses, $\alpha_{jk}$, were empirically estimated from reductions in a species' growth rate across conspecific or heterospecific densities ranging from 0 to >1000 individuals at 12 and 28°C (7) (Fig. S2). Turion production and germination functions (eqs. 4 & 5) were fitted using values from the literature (4, 5). All parameter values are shown in table S1.

Following model selection, our final model (eq. 1 in the main text) was able to predict the observed *per capita* growth rates of both *S. polyrhiza* and *L. minor* with good accuracy across a range of temperatures (predicted  observed $R^2 = 0.76, 0.80$, for each species, respectively) (Fig. S3) (2).

**Detailed Methods for MaxEnt Ecological Niche Models.** Spatial point occurrence records for each species were downloaded using the *rgbif* package (8). This package facilitates remotely accessing data from the Global Biodiversity Information Facility database (GBIF; http://gbif.org; accessed February 2019) using the R language (9). After downloading all georeferenced records for *Lemna minor* (193,395 records, https://doi.org/10.15468/dl.wpisn8) and *Spirodela polyrhiza* (83,531 records; https://doi.org/10.15468/dl.2pixjr). We cleaned the data using the R package *CoordinateCleaner* (10), which flags and removes problematic entries. Our specific omission criteria included:

1. Points falling within a 10 km radius of a country's capital city.
2. Points falling within a 1 km radius of a country's centroid coordinate.
3. Points possessing identical longitude and latitude values.
4. Points falling within a 1° radius of the GBIF headquarters in Copenhagen, Denmark.
5. Points falling within a 1 km radius of biodiversity institutions such as herbaria.
6. Points determined to be in the ocean.
7. Points having at least one coordinate exactly equal to zero degrees.
8. Points collected prior to 1900.
9. Points outside of the study regions of interest (Mexico, USA, Canada, United Kingdom, Ireland, and northern continental Europe). Note that we also retained all global records to create fig. 4 in the main text.

We amended these occurrence records with gridded survey records from the UK and Ireland provided by the Botanical Society of Britain and Ireland (BSBI; ver. February 2019) (11). Point locations for these data represent the centroids of 10 km$^2$ grid cells in which a species was observed. Despite the gridded nature of these observations, the sampling scheme is far more thorough than data obtained through GBIF, and are trusted to represent the true spatial extent of the study species across the UK and Ireland. Given the high density of occurrence records in our study regions, we made the assumption that the distribution of occurrence records reflected the true geographic distributions of our study species. We note that while high-latitude observations may have been missed due to a lack of sampling effort (particularly in N. America), the observed distributions of points align closely with distributional accounts in the literature (1, 12).

Because ecological niche models can be sensitive to the effects of spatially-clustered and potentially duplicated occurrence records (13), we randomly sampled one individual record falling within cells arranged in a gridded overlay. For our regional occurrence datasets, the cell sizes were 1.3° x 1.3°, 1° x 1°, and 0.3° x 0.3° for the N. America, N. EU, and UK datasets, respectively. We generated a fourth dataset by combining the unfiltered observations in all three regions, and then then spatially filtering these points by sampling from 1.5° x 1.5° grid cells. The number of records remaining in each regional dataset before and after correcting for spatial sampling bias is summarized in table S1.

Two different sets of environmental covariates were used to fit statistical niche models. The first included 12 bioclimatic variables downloaded from the WorldClim database (14) at 2.5 arcmin resolution. We selected these variables based on their hypothesized or known contributions to the growth of aquatic plants. Our second set of covariates includes only the BIO1 (average annual temperature) and BIO7 (annual temperature amplitude) measurements, which are the same variables used in our mechanistic invasion model predictions (eq. 1). Summaries of these covariates and the models in which they were used can be found in table S2. These bioclimatic variables were then clipped to the extents of our study regions. We conducted principal component analyses (PCA) on these environmental variables to obtain the first two principal component axes (Fig. S5) and tested how the first environmental principal component varied across latitude in each region using linear regression.

We used the MaxEnt software (version 3.4) (15) implemented within the R package *ENMeval* (16) to create niche models for our two duckweed species. MaxEnt is a statistical niche modeling framework that uses environmental covariates extracted from spatial occurrence data to predict a species' habitat suitability relative to a randomly-sampled environmental background (17). As a presence-only method, MaxEnt can use the type of aggregated occurrence records stored on the GBIF, and despite not permitting information on species' absences, performs favorably in comparisons with other statistical niche modeling approaches (18). For each of our species (*L. minor* and *S. polyrhiza*) and set of covariates (2 or 12 BIOCLIM variables), we generated individual MaxEnt models for each of our three study regions, as well as a fourth, combined region. Environmental background points were generated within each of these regions by randomly sampling between 10,000 (UK & Ireland) and 30,000 (combined regions) random points that did not coincide with a species' observation record.

We employed two strategies to avoid overfitting the MaxEnt models. First, we generated four nested, spatial partitions of the presence and background data, which were then used for 4-fold cross validation across spatially-segregated training and test datasets (19). Second, we fit our models using a range of regularization parameters (1.5 through 6). Higher values of the regularization parameters result in more general, smoother model predictions (17). MaxEnt models were fit with combinations of linear, quadratic, hinge, and product feature classes using *ENMeval* (16) with clamping enabled. Best-fit MaxEnt models were selected using a pluralistic approach to improve precision while controlling for overfitting. We favored models with a combination of low AIC values (20), average 10% threshold omission rates closest to 0.1, and higher regularization parameters. MaxEnt habitat suitability thresholds were estimated using a 10% omission rate criterion. One-sided binomial tests of omission were used to determine model discriminatory performance. This test estimates the probability that a given MaxEnt or mechanistic invasion model's predictions are significantly better than what would be obtained by chance alone (21).

**Estimating Range Limits.** We used an inverse regression approach (22) to estimate the latitudinal limits of *S. polyrhiza* from model outputs. For each study region, we extracted both invasion and MaxEnt model outputs from cells containing an observation record. After fitting a linear regression model to these points, we estimated these models' $x$-intercepts and their inverse 95% confidence intervals, which represent predictions for the latitudinal limit of *S. polyrhiza*. For our invasion model, this limit is defined as the latitude at which the low-density growth rate, $\bar{r}_{\text{inv}}$, equals zero. For MaxEnt models, it was defined as the latitude at which the cloglog occurrence probability, $p_{\text{occ}}$, equalled the 10% omission threshold, $\tau$. While choosing such a binary presence/absence threshold is subjective, it is necessary for quantifying a range boundary, and we found that a 10% omission threshold resulted in good approximations of the species' observed distributions. Since $\bar{r}_{\text{inv}}$ and $p_{\text{occ}}$ can be nonlinear over latitude, we performed regressions only on data points near the maximum latitude in each region. These subsets were selected where the latitude-output response appeared linear and included 138, 244, and 73 location records for the UK + Ire., N. America, and N. EU study regions, respectively.

**Partitioning Coexistence Mechanisms.** Modern coexistence theory (MCT) provides a conceptual and analytical framework for partitioning the effects of various fluctuation-dependent and fluctuation-independent mechanisms contributing to coexistence (23). As explained in the main text, stable coexistence hinges on satisfying the reciprocal invasibility criterion (i.e., $\bar{r}_{j\setminus\text{inv}} > 0$ for all species in a community) (24, 25). The relative strength of each mechanism is then obtained by calculating invader-resident differences. A standard, non-spatial MCT partitioning uses Taylor approximations to decompose the invasion growth rate into three primary components: (1) the *fluctuation-independent growth rate*, which is the growth rate in the absence of fluctuations in competition, and includes the effects of intrinsic growth and negative density-dependence arising from niche differences, (2) *relative nonlinearity in competition*, which measures the differential impacts of nonlinear averaging (and thereby Jensen's inequality) on species' nonlinear responses to competition, and (3) the *storage effect*, which measures the buffering of population losses in harsh times relative to large gains in favorable times, and is a form of temporal niche partitioning (26, 27). Importantly, the latter two mechanisms can only occur when environments or resources fluctuate through time.

We used a recently-developed computational method (28) for partitioning each species' invasion and resident growth rates, $\bar{r}_{j\setminus\text{inv}}$ and $\bar{r}_{j\setminus\text{res}}$, into terms reflecting the additive contributions of fluctuation-dependent and independent coexistence mechanisms (23, 29). This method, while accurate, results in slightly different, though arguably more interpretable results than the classic small-variance approximations used for partitioning in the classical MCT approach (27, 29). We briefly outline this approach below, but refer interested readers to the original literature for a more thorough overview (28, 30).

Our invasion models contain three variable quantities that can affect each species' invasion and resident growth rates: conspecific densities ($N_j(t)$), heterospecific densities ($N_k(t), k \neq j$), and temperature, ($T(t)$). Species' long-term average invasion and resident growth rates can be written as

$$\bar{r}_{j\backslash\text{inv}} = \frac{1}{m}\sum_{v=1}^{m} r_j\big(T(t_s), N_k(t_s)\big)$$
$$\bar{r}_{j\backslash\text{res}} = \frac{1}{m}\sum_{v=1}^{m} r_j\big(T(t_s), N_j(t_s)\big), \quad \text{[S1]}$$

where $r_j(T, N_k) = (N_j + S_j)^{-1}(dN_j/dt + dS_j/dt)$, and $t_v(v = 1, ..., m)$ are finely-spaced time points stretching over 365 total days. Following (28), we can partition these average *per capita* rates such that

$$\bar{r}_j = \varepsilon_j^* + \varepsilon_j' + \bar{\varepsilon}_j^T + \bar{\varepsilon}_j^{N_k} + \bar{\varepsilon}_j^{(T\#N_k)} + \bar{\varepsilon}_j^{(TN_k)}, \quad \text{[S2]}$$

where $k = j$ for the species in its resident state, and $k \neq j$ in its invasion state. The $\varepsilon_j$ terms are defined as

| | | |
|---|---|---|
| $\varepsilon_j^*$ | $= r_j(\bar{T}, \bar{N}_k^*)$, | the fluctuation-free growth rate |
| $\varepsilon_j'$ | $= r_j(\bar{T}, \bar{N}_k) - \varepsilon_j^*$, | the effect of fluctuation-driven change in mean $N_k$ |
| $\bar{\varepsilon}_j^T$ | $= \frac{1}{m}\sum_{v=1}^{m} r_j(\bar{T}, N_k(t_v)) - [\varepsilon_j^* + \varepsilon_j']$, | the main effect of variation in temperature |
| $\bar{\varepsilon}_j^{N_k}$ | $= \frac{1}{m}\sum_{v=1}^{m} r_j(T(t_v), \bar{N}_k) - [\varepsilon_j^* + \varepsilon_j']$, | the main effect of variation in competitor density |
| $\bar{\varepsilon}_j^{TN_k}$ | $= \frac{1}{m}\sum_{v=1}^{m} r_j(T(t_v), N_k(t_v)) - [\varepsilon_j^* + \varepsilon_j' + \bar{\varepsilon}_j^T + \bar{\varepsilon}_j^{N_k}]$, | the interaction of $T$ and $N_k$ variation |
| $\bar{\varepsilon}_j^{(T\#N_k)}$ | $= \frac{1}{m^2}\sum_{v=1}^{m}\sum_{w=1}^{m} r_j(T(t_v), N_k(t_w)) - [\varepsilon_j^* + \varepsilon_j' + \bar{\varepsilon}_j^T + \bar{\varepsilon}_j^{N_k}]$, | the independent variation component of $\bar{\varepsilon}_j^{TN_k}$ |
| $\bar{\varepsilon}_j^{(TN_k)}$ | $= \bar{\varepsilon}_j^{TN_k} - \bar{\varepsilon}_j^{(T\#N_k)}$, | the covariance component of $\bar{\varepsilon}_j^{TN_k}$, |

where $\bar{T}$ and $\bar{N}_k$ are arithmetic averages taken over the final 365 days of the simulation. $\bar{N}_k^*$ represents the average value of $N_k$ when temperatures do not fluctuate, which in our model, are point equilibria and therefore constant over time. Next, like in the analytical treatment of modern coexistence theory, we performed invader-resident comparisons to assess the relative extent to which each of the terms above benefits or harms the invading species. For example, we can define $\Delta_j^T = \bar{\varepsilon}_{j\backslash\text{inv}}^T - \bar{\varepsilon}_{k\backslash\text{res}}^T$ $(j \neq k)$ as the relative (dis)advantage experienced by an invader owing solely to the species' varying responses to temperature fluctuations. This term and $\Delta_j^{N_k}$ measure the relative nonlinearity in species' responses to temperatures and competitor densities. Likewise, $\Delta_j^{(TN_k)} = \bar{\varepsilon}_{j\backslash\text{inv}}^{(TN_k)} - \bar{\varepsilon}_{k\backslash\text{res}}^{(TN_k)}$ represents the contribution of covariance between the environment ($T$) and competitive factor ($N_k$) to the invader's growth rate. This term quantifies the coexistence mechanism called the temporal storage effect (27). For our analysis, we set the MCT invader-resident comparison quotients, $q_{ir}$, to 1, since they cannot be uniquely defined for all of our environmental states (28, 30), and are very close to 1 when unique solutions exist (2).

We calculated the growth components for each species in its invader state at each point across a grid of average temperatures ranging from -10 to 37 °C with amplitudes ranging from 0 to 33 °C. These values span the natural range of temperature spaces across Earth. As described in the main text, we first simulated the dynamics of the resident species for 10 years, checking that its long-term growth rate was approximately zero. We used the final year of resident densities and temperatures to calculate the growth rate of an invader at each time step, which was then geometrically-averaged to obtain $\bar{r}_{j\backslash\text{inv}}$. Partitioning then proceeded as described above and in (28). To check our results, we verified that the following relations held for each species in its invader state:

$$\begin{aligned}\bar{r}_{j\backslash\text{inv}} &= \varepsilon_j^* + \varepsilon_j' + \bar{\varepsilon}_j^T + \bar{\varepsilon}_j^{N_k} + \bar{\varepsilon}_j^{(T\#N_k)} + \bar{\varepsilon}_j^{(TN_k)} \\ &\approx \bar{r}_{j\backslash\text{inv}} - \bar{r}_{k\backslash\text{res}} \\ &\approx \Delta_j^* + \Delta_j' + \Delta_j^T + \Delta_j^{N_k} + \Delta_j^{(T\#N_k)} + \Delta_j^{(TN_k)}, \quad (j \neq k).\end{aligned} \quad \text{[S3]}$$

We plotted our results as a heatmap over our 2-D temperature-amplitude grid, onto which we overlaid spatially-thinned global datasets of *S. polyrhiza* or *L. minor* observation records (figs. 5 and S4) obtained from GBIF and BSBI. We also overlaid the zero-growth isoclines for each species.

**Estimating stabilizing and equalizing components.** We also calculated the stabilizing and equalizing components of the $\Delta_j^i$ partitions in eq. S3 (28, 31). Here, stabilizing components $\overline{\Delta^i}$ represent the average contribution of a particular coexistence mechanism $i$ to competitors' invasion growth rates and equals the arithmetic average across species of a particular $\Delta^i$. Positive stabilization will help both species increase when rare. Equalizing effects arise when a particular coexistence mechanism reduces fitness differences between an invader and resident and are equal to $\Delta_j^i - \overline{\Delta^i}$. Thus, a particular component of the growth rate partition (e.g., the storage effect, $\Delta_j^{(TN_k)}$) can result in any combination of stabilization and equalization terms depending the direction and relative magnitude of its actions on both species. Summed across a species, the terms will equal the species' invasion growth rate. Note that although this stabilization value can be positive even when an individual contribution of $\Delta_j^i$ is negative, our usage of $\overline{\Delta^T}$ was positive only when both $\Delta_j^i$'s were greater than zero.

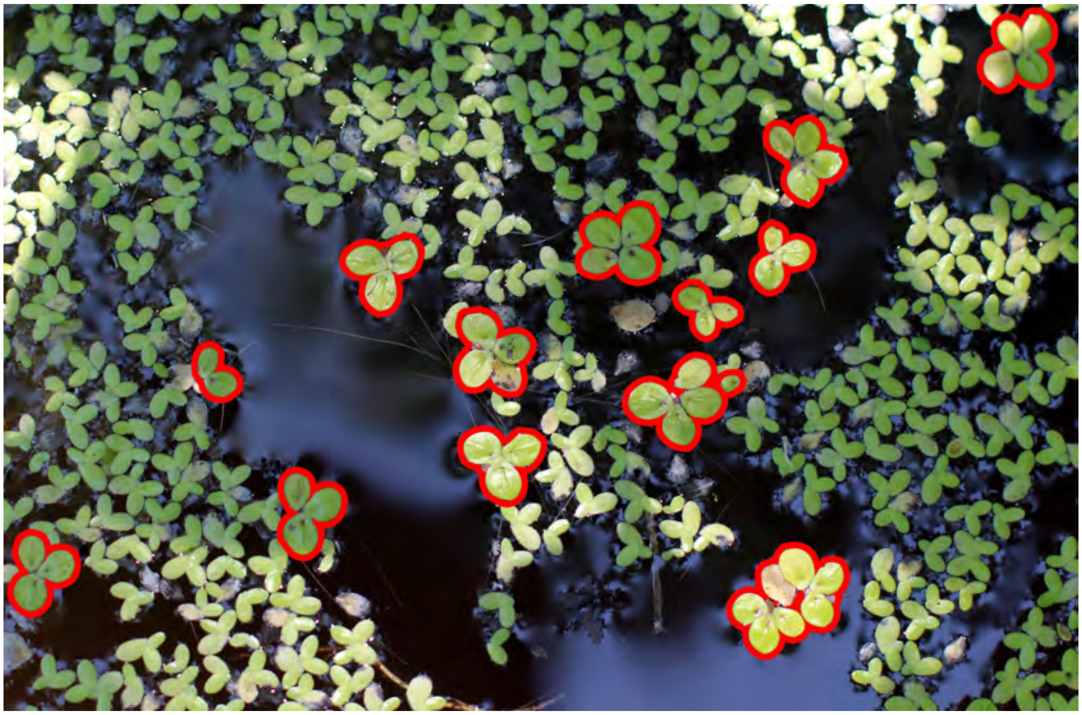

**Fig. S1.** A typical mixed community of *Lemna minor* and *Spirodela polyrhiza* (outlined in red).

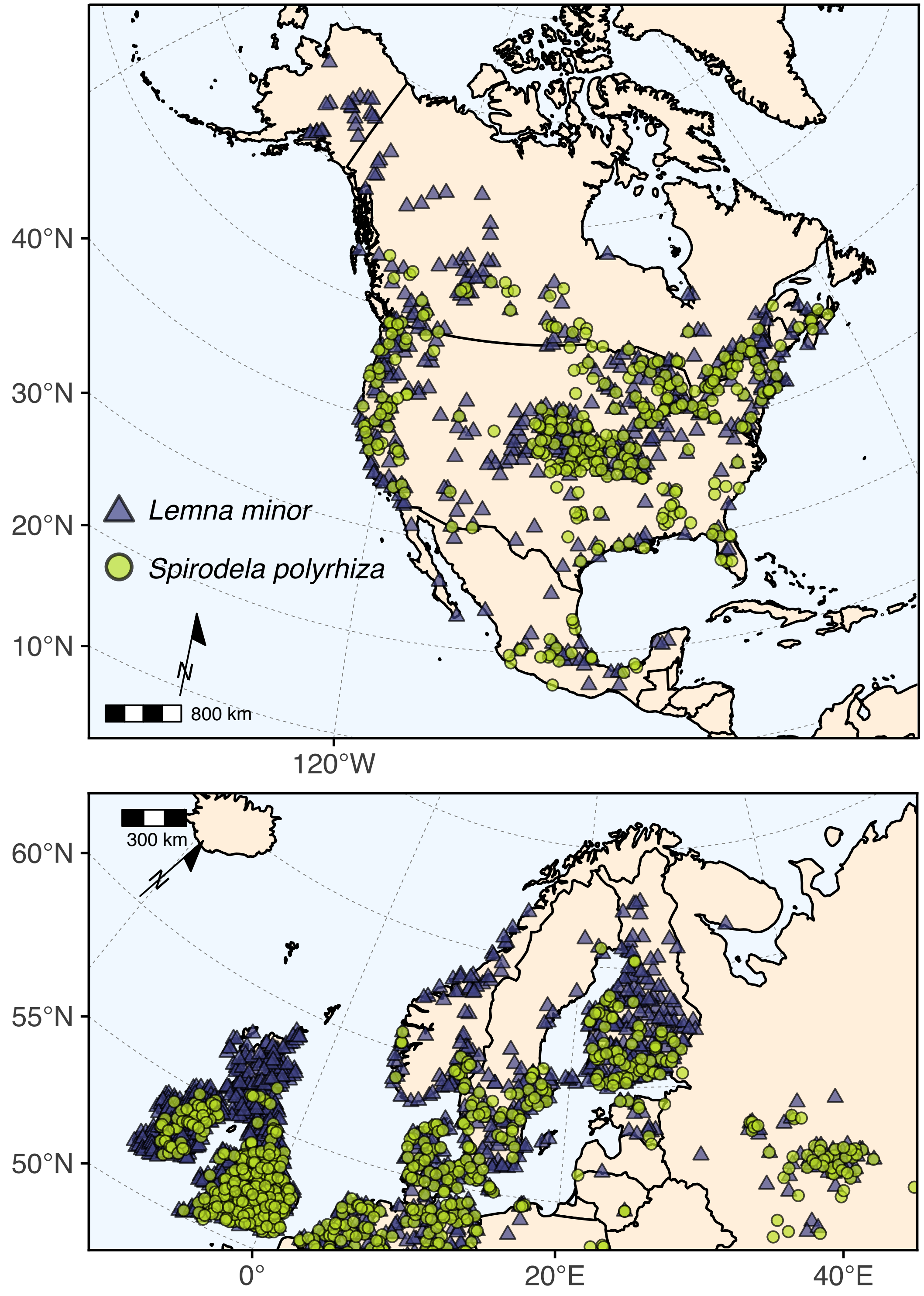


**Fig. S2.** Maps of location records for *Lemna minor* and *Spirodela polyrhiza* for North America (CA, USA, MX) and Northern Europe. Note the greater northern range limits for *L. minor* relative to *S. polyrhiza*. Points have been spatially-thinned for easier viewing.

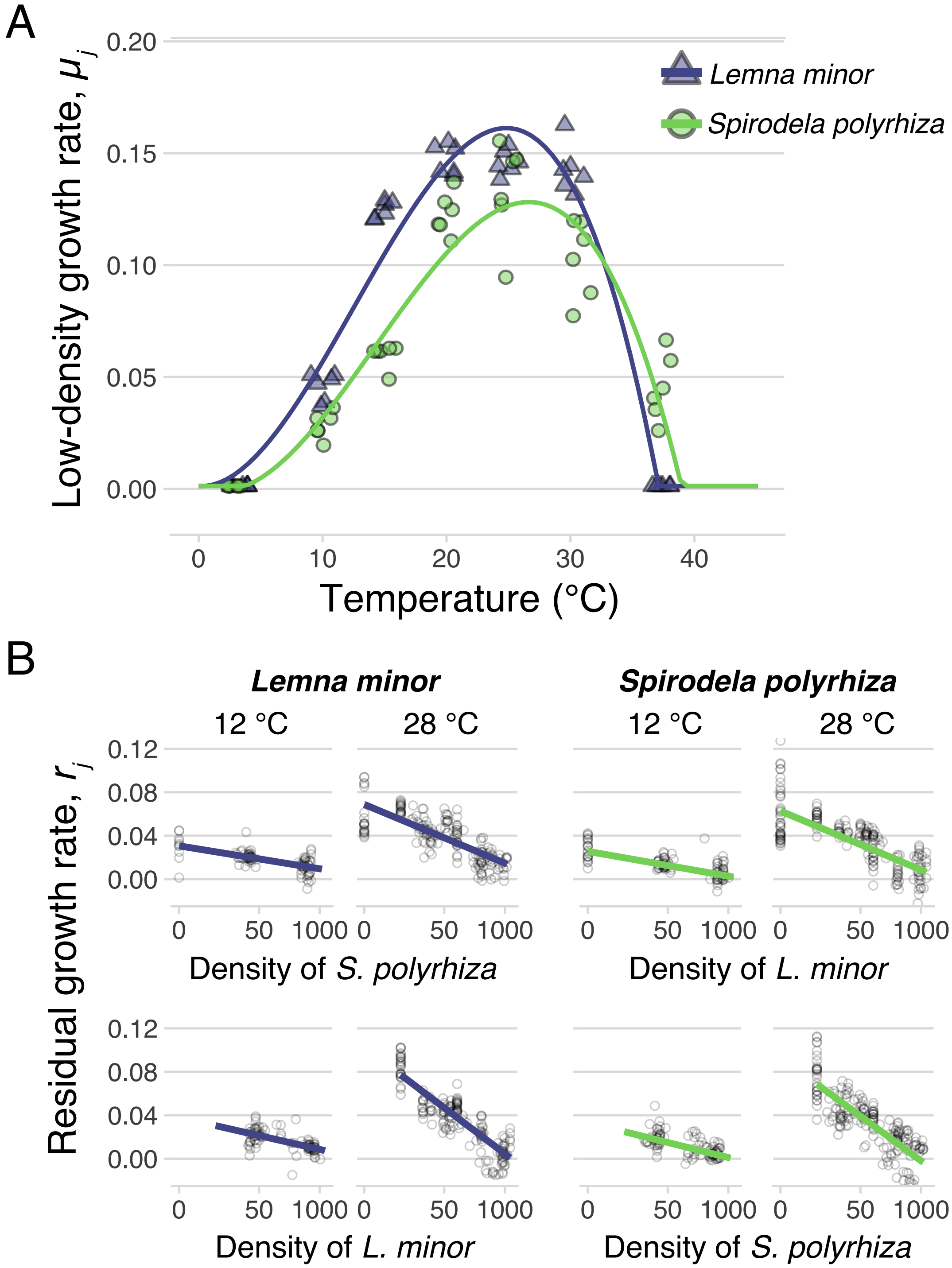


**Fig. S3.** (A) Empirically-measured thermal growth maxima, $\mu_j(T)$ (per day), for *L. minor* and *S. polyrhiza* (data from (2)). Curves were fit using equation 2. (B) Partial residual plots showing the effects of ambient temperature on interspecific (top) and intraspecific (bottom) negative density dependence for each species. Points are partial residuals of growth rates $r_j$ (per day) from experimental cultures (2). Regression lines show the conditional effects of heterospecific and conspecific densities, controlling for the impacts of the other species.

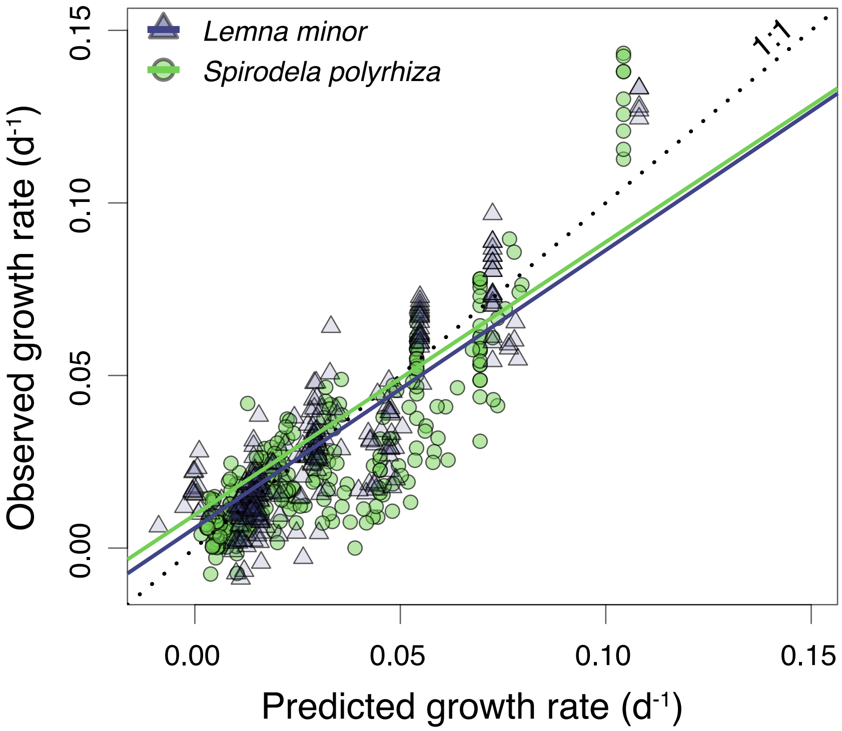


**Fig. S4.** Relationship between model-predicted and observed growth rates for *L. minor* and *S. polyrhiza*. Model predictions use equation 1 with parameters estimated from growth experiments. Observed values are from measurements taken in competition assays under fluctuating temperatures (see (2) for details). State variables used to generate predictions were temperature, conspecific density, and heterospecific density. Dotted line denotes perfect predictive accuracy. Regression lines for each species are also displayed.

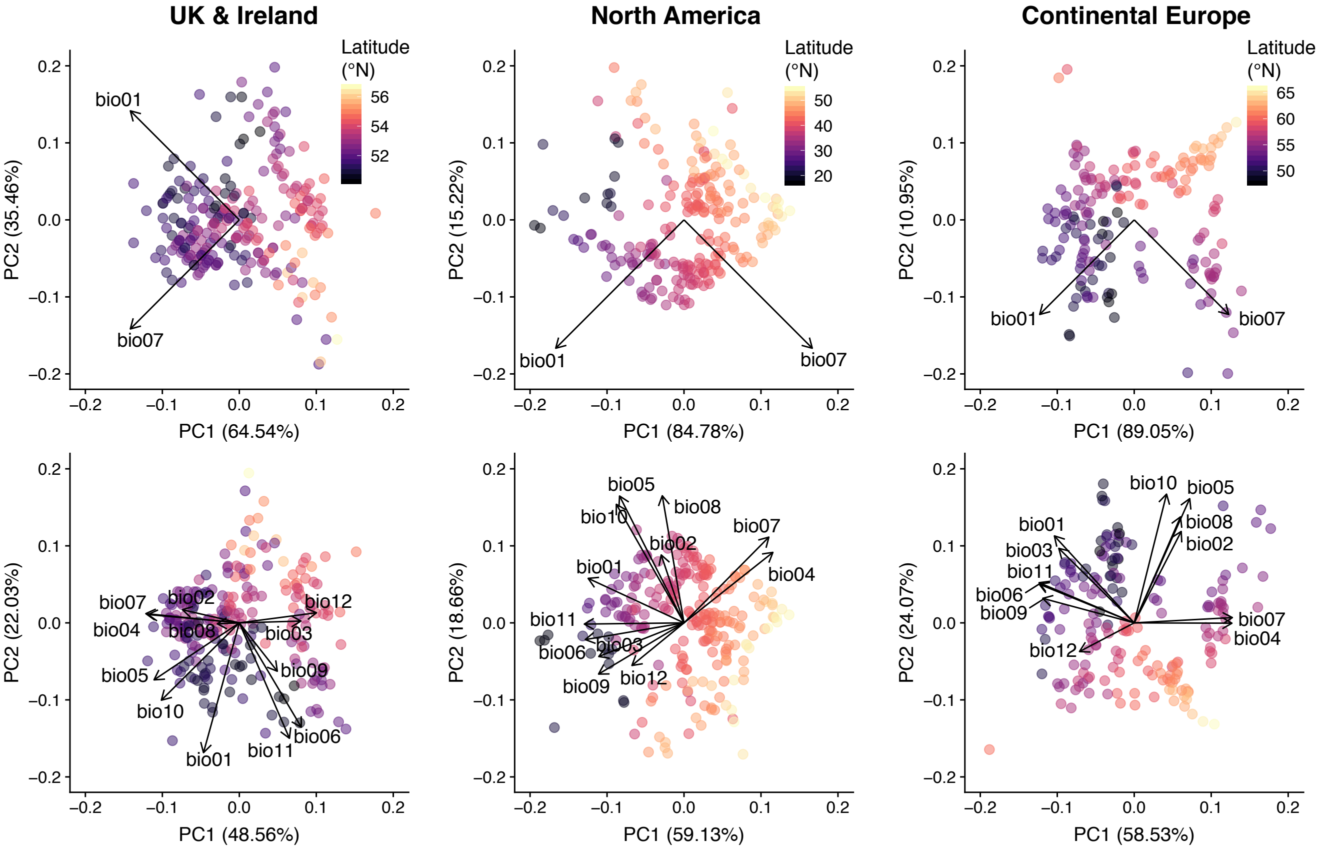


**Fig. S5.** Principal components biplots showing first two PCA axes for environmental covariates for each study region. Points represent observations of *S. polyrhiza*, and are shaded by latitude, and arrows denote loading of each variable. The top row shows results for mean temperature (bio01) and temperature amplitude (bio07) variables — used to fit invasion models and 2-variable MaxEnt models. Second row contains results for variables used in the 12-variable MaxEnt model (bio1-bio12). Descriptions of these variables can be found at http://www.worldclim.org/bioclim. Values on axis labels denote percentage of variance explained by the first and second PC axes.

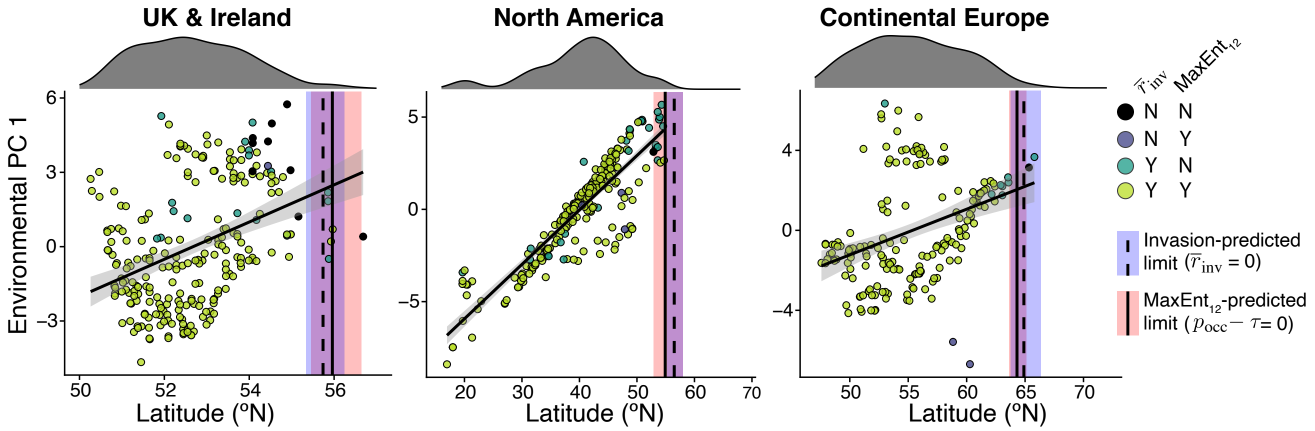


**Fig. S6.** Latitudinal limit predictions for invasion model and 12-variable MaxEnt model. Points denote *S. polyrhiza* observation records, colored by model classification results, and vertical lines signify the estimated latitudinal limits for each model type and region. Histograms above each plot show the latitudinal dispersion of occurrence records after spatial thinning.

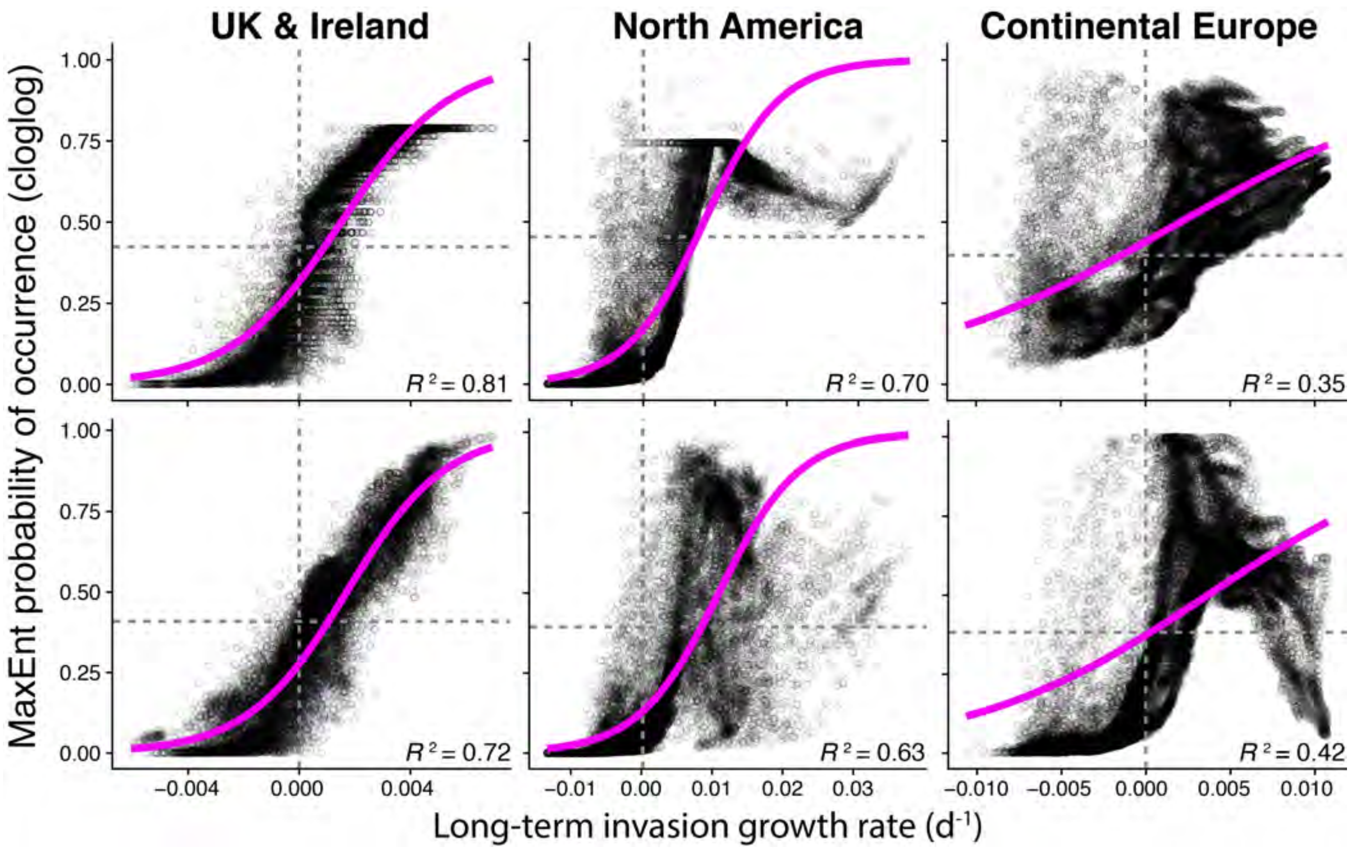


**Fig. S7.** Relationships between invasion model-predicted growth rates, $\bar{r}_{\rm inv}$, and MaxEnt-predicted occurrence probabilities for *S. polyrhiza*. Points represent values for each 2.5 arcmin grid cell. Horizontal and vertical dashed lines indicate the MaxEnt presence/absence threshold and low-density growth thresholds, respectively. Colored lines show results from beta regression models. The top and bottom rows contain data from the 2 and 12-variable MaxEnt models, respectively.

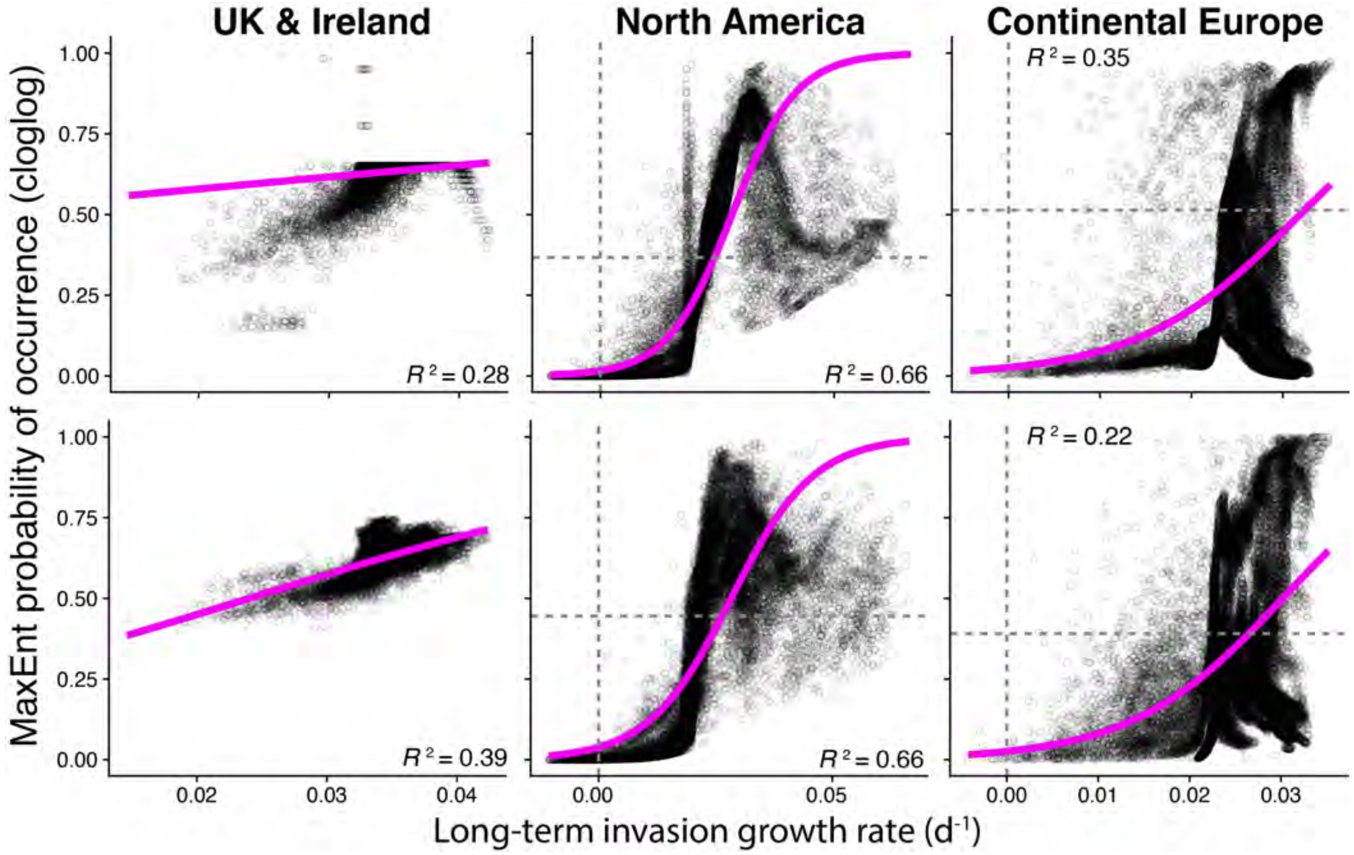


**Fig. S8.** Relationships between invasion model-predicted growth rates, $\bar{r}_{\mathrm{inv}}$, and MaxEnt-predicted occurrence probabilities for *L. minor.* Points represent values for each 2.5 arcmin grid cell. Horizontal and vertical dashed lines indicate the MaxEnt presence/absence threshold and low-density growth thresholds, respectively. Colored lines show results from beta regression models. The top and bottom rows contain data from the 2 and 12-variable MaxEnt models, respectively.

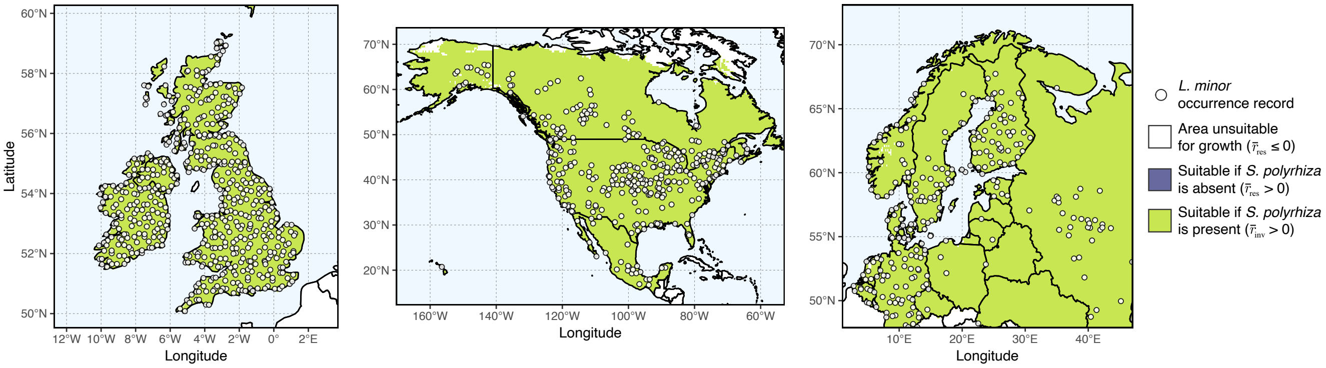


**Fig. S9.** Range predictions for *L. minor* from the competition model (eq. 1) projected across geographic space. Binary outcomes show areas of predicted population persistence satisfying the invasion criterion ($\bar{r} > 0$). The predicted latitudinal limits of *L. minor* do not change if competition from resident *S. polyrhiza* is accounted for.

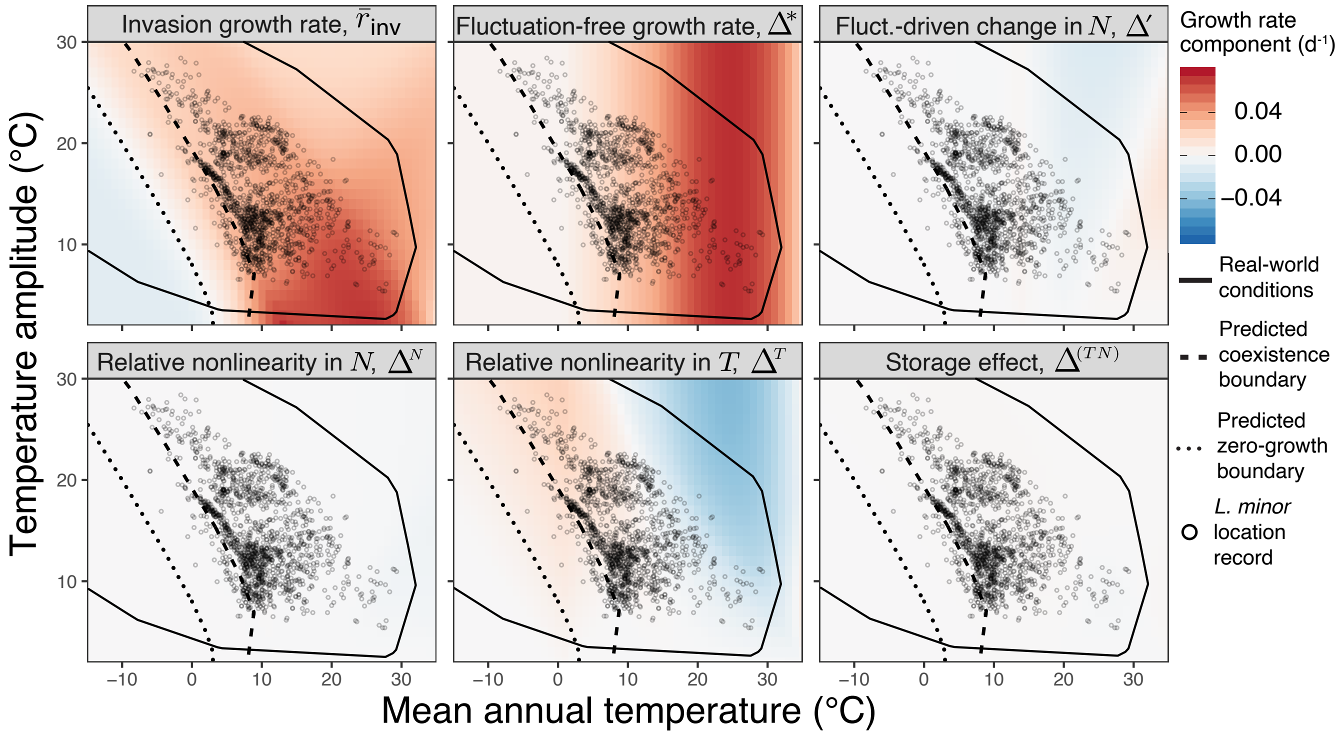


**Fig. S10.** Long-term invasion-growth rate partitioning for *Lemna minor*. The upper-left panel displays the overall growth rate, $\bar{r}_{\text{inv}}$, and the following panels reflect the contributions of various coexistence-promoting mechanisms from equation 3. Points represent thinned global occurrence records for *L. minor*. Both coexistence boundaries and zero-growth boundaries are shown.

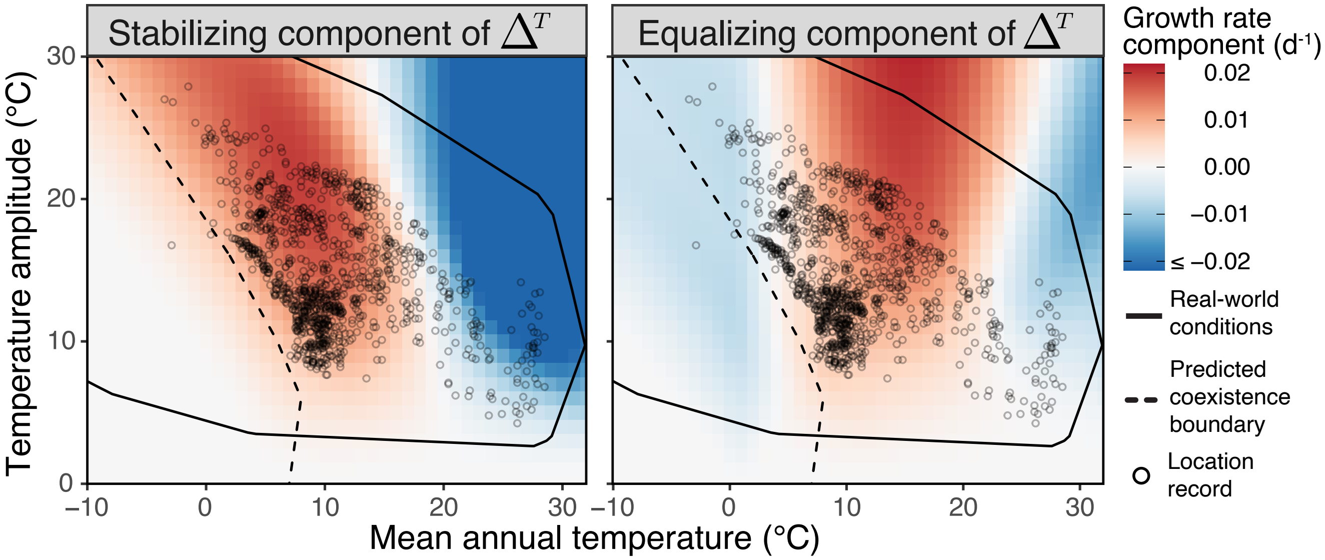


**Fig. S11.** Stabilizing and equalizing components of $\Delta^T$ for *Spirodela polyrhiza*. Stabilization in $\Delta^T$ occurs when relative nonlinearity in thermal responses facilitates both species' invasion growth rates. Equalization in $\Delta^T$ occurs when relative nonlinearity in thermal responses reduces the difference in invasion growth rates, favoring the species with the lower growth rate. Stabilizing components of $\Delta^T$ are the same for both species, while equalizing components for *L. minor* are opposite those of *S. polyrhiza*. Points represent thinned global occurrence records for *S. polyrhiza*.

**Table S1. Parameter values used for simulating mechanistic niche models. See (2) for estimation procedures.**

| Parameter | Description | Species | Value (± 95% CI) |
|---|---|---|---|
| $T_{\mathrm{max},j}$ | Maximum growth temperature (℃) | *S. polyrhiza* | $38.9$ $(0.4)$ |
| | | *L. minor* | $37.0$ $(0.7)$ |
| $T_{\mathrm{min},j}$ | Minimum growth temperature (℃) | *S. polyrhiza* | $3.8$ $(1.6)$ |
| | | *L. minor* | $0.5$ $(1.1)$ |
| $c_j$ | Scaling constant for thermal growth model | *S. polyrhiza* | $1.7 \times 10^{-5}$ $(1.8 \times 10^{-6})$ |
| | | *L. minor* | $2.2 \times 10^{-5}$ $(1.3 \times 10^{-6})$ |
| $T_{\mathrm{d},j}$ | Temperature at which 50% of growth is devoted to turions (℃) | *S. polyrhiza* | $15$ |
| | | *L. minor* | n/a |
| $T_{\mathrm{g},j}$ | Temperature at which 50% of turions germinate at 20 days (℃) | *S. polyrhiza* | $25$ |
| | | *L. minor* | n/a |
| $\alpha_{jj}(\overline{T})$ | Intraspecific competition parameter (at 20℃) | *S. polyrhiza* | $0.1069$ $(0.016)$ |
| | | *L. minor* | $0.1005$ $(0.015)$ |
| $\alpha_{jk}(\overline{T})$ | Interspecific competition parameter (at 20℃) | *S. polyrhiza* | $0.0646$ $(0.009)$ |
| | | *L. minor* | $0.0591$ $(0.010)$ |
| $\psi_{jj}$ | Effect of 1℃ temperature change on $\alpha_{jj}$ | *S. polyrhiza* | $6.2 \times 10^{-3}$ $(1.9 \times 10^{-3})$ |
| | | *L. minor* | $6.3 \times 10^{-3}$ $(1.7 \times 10^{-3})$ |
| $\psi_{jk}$ | Effect of 1℃ temperature change on $\alpha_{jk}$ | *S. polyrhiza* | $3.1 \times 10^{-3}$ $(1.1 \times 10^{-3})$ |
| | | *L. minor* | $3.0 \times 10^{-3}$ $(1.1 \times 10^{-3})$ |
| $m_j$ | Species' average *per capita* mortality rate $(d^{-}1)$ | *S. polyrhiza* | $0.0134$ $(7.4 \times 10^{-4})$ |
| | | *L. minor* | $0.0107$ $(6.8 \times 10^{-4})$ |

**Table S2. Numbers of cleaned observation records for each of the study species both before and after spatial bias correction via grid filtering. "Combined" row represents points aggregated from the three study regions.**

| Species | Region | Before spatial filter | After spatial filter |
|---|---:|---:|---:|
| *L. minor* | World | 153,575 | 445 |
| | N. America | 1,402 | 340 |
| | N. EU | 104,050 | 272 |
| | UK + Ireland | 22,470 | 546 |
| | Combined | 127,922 | 750 |
| *S. polyrhiza* | World | 69,989 | 315 |
| | N. America | 897 | 225 |
| | N. EU | 58,906 | 192 |
| | UK + Ireland | 2,266 | 250 |
| | Combined | 62,069 | 516 |

**Table S3. Bioclimatic variables used to fit ecological niche models.**

| Name | Description | $\bar{r}_{\mathrm{inv}}$ | $\mathrm{MaxEnt}_2$ | $\mathrm{MaxEnt}_{12}$ |
|---|---|---|---|---|
| BIO1 | Annual mean temperature | Y | Y | Y |
| BIO2 | Mean of monthly diurnal temperature range | | | Y |
| BIO3 | Isothermality (BIO2/BIO7) | | | Y |
| BIO4 | Temperature seasonality (annual standard deviation) | | | Y |
| BIO5 | Max temperature of warmest month | | | Y |
| BIO6 | Min temperature of coldest month | | | Y |
| BIO7 | Annual temperature amplitude | Y | Y | Y |
| BIO8 | Mean temperature of wettest quarter | | | Y |
| BIO9 | Mean temperature of driest quarter | | | Y |
| BIO10 | Mean temperature of warmest quarter | | | Y |
| BIO11 | Mean Temperature of coldest quarter | | | Y |
| BIO12 | Annual precipitation | | | Y |

**Table S4. Fit statistics for 2- and 12-covariate MaxEnt models. Features include hinge (H), linear (L), quadratic (Q), and polynomial (P) functions, or any combination of the four. RM indicates the regularization multiplier. Area-under-curve (AUC) values are calculated by averaging across 4 independently subsetted evaluation datasets. △AUC show average differences between individual calibration and evaluation AUC values. Omission rate (OR) metrics can be compared to theoretical expectations of omission rates of 0 ($OR_{MTP}$) and 0.1 ($OR_{10\%}$). Omission rates equal to or lower than their theoretical expectations and △AUC values closer to zero signify lower overfitting.**

| Species | Region | Covariates | Features | RM | AUC | △AUC | $OR_{MTP}$ | $OR_{10\%}$ | Parameters |
|---|---|---|---|---|---|---|---|---|---|
| *S. polyrhiza* | UK & Ireland | 2 | LQHP | 3.5 | 0.74 | 0.01 | 0.007 | 0.098 | 6 |
| | | 12 | LQH | 4 | 0.74 | 0.02 | 0.011 | 0.102 | 16 |
| | N. America | 2 | LQ | 1.5 | 0.77 | 0.02 | 0.002 | 0.093 | 3 |
| | | 12 | LQ | 1.5 | 0.81 | 0.02 | 0.012 | 0.116 | 12 |
| | N. Europe | 2 | H | 4 | 0.71 | 0.03 | 0.003 | 0.091 | 18 |
| | | 12 | LQ | 1.5 | 0.80 | 0.02 | 0.001 | 0.086 | 14 |
| *L. minor* | UK & Ireland | 2 | H | 1.5 | 0.67 | 0.03 | 0.003 | 0.090 | 37 |
| | | 12 | H | 3 | 0.76 | 0.02 | 0.002 | 0.090 | 35 |
| | N. America | 2 | L | 4 | 0.65 | 0.01 | 0.004 | 0.101 | 2 |
| | | 12 | L | 3 | 0.66 | 0.02 | 0.003 | 0.101 | 5 |
| | N. Europe | 2 | L | 2 | 0.73 | 0.01 | 0.001 | 0.109 | 2 |
| | | 12 | H | 3 | 0.76 | 0.02 | 0.002 | 0.090 | 35 |

**Table S5. Comparison of *S. polyrhiza*'s observed northern latitudinal limits, $\overline{L}_{\max}$ (° N), with niche model-estimated latitudinal limits, $\hat{L}_{\max}$. Values in parentheses denote 95% confidence intervals. True positive rates (TPR) and binomial omission test results (*p*-values) are used to assess model model fit to observation records. For model definitions, see Materials and Methods section.**

| Region | $\overline{L}_{\max}$ | Model | $\hat{L}_{\max}$ | TPR |
|---|---|---|---|---|
| UK & Ireland | (53.7, 56.3) | $\bar{r}_{\mathrm{res}}$ | (65.0, 68.5) | $1.00^{\mathrm{n.s.}}$ |
| | | $\bar{r}_{\mathrm{inv}}$ | (55.3, 56.2) | 0.95** |
| | | $\mathrm{MaxEnt}_2$ | (55.4, 56.7) | 0.90** |
| | | $\mathrm{MaxEnt}_{12}$ | (55.2, 56.5) | 0.89** |
| N. America | (48.6, 55.0) | $\bar{r}_{\mathrm{res}}$ | (62.7, 66.0) | $1.00^{\mathrm{n.s.}}$ |
| | | $\bar{r}_{\mathrm{inv}}$ | (55.5, 58.3) | 0.98** |
| | | $\mathrm{ME}_2$ | (51.9, 54.7) | 0.91** |
| | | $\mathrm{ME}_{12}$ | (53.4, 60.0) | 0.90** |
| N. Europe | (61.8, 66.0) | $\bar{r}_{\mathrm{res}}$ | (77.2, 80.7) | 1.00* |
| | | $\bar{r}_{\mathrm{inv}}$ | (63.8, 66.5) | 0.98** |
| | | $\mathrm{ME}_2$ | (62.6, 63.4) | 0.90** |
| | | $\mathrm{ME}_{12}$ | (63.7, 65.2) | 0.95** |

$^{\mathrm{n.s.}}p > 0.05; {}^{*}p < 0.005; {}^{**}p < 0.001$
